# Model based acoustic gaze inference for behaviour in the field, array design and honest uncertainty estimation

**DOI:** 10.64898/2026.11.19.767943

**Authors:** Thejasvi Beleyur

**Affiliations:** Active Sensing Collectives Lab, Dept. of Biology, University of Konstanz, 78464, Konstanz, DE

## Abstract

Active sensing animals like echolocating bats and porpoises modulate their sensory volumes in a directed manner by altering various aspects (aim, beam width, source-level) of their calls or clicks, termed ‘acoustic gaze’. Studying acoustic gaze requires inferring the beam parameters from a limited set of sensors, which is however a non-trivial task. To aid inference, studies apply a series of experimental constraints and data curation practices (e.g. animals always facing towards the array, only calls with peaks within X degrees of the centre analysed). Particularly for field studies, these constraints drastically reduce the volume of data, and remove important spatio-temporal context. Here I present a model-based Bayesian paradigm to perform acoustic gaze inference that releases the need for arbitrary constraints and curation practices such as animal flight direction or aim with three computational demonstrations 1) recovering acoustic gaze from a free-flying animal without constraints 2) post-hoc calibration of previously published array geometries that expand the spatial range of analysable calls and 3) a priori calibration of a proposed array design to avoid wasted data collection. The model-based paradigm expands the conditions under which acoustic gaze inference can be done, while also providing a principled approach to honestly estimate the uncertainty in estimated parameters.

## I. INTRODUCTION

Active sensing animals are able to control their sensory volumes by jointly modulating the source level, directionality and acoustic beam aim of their emitted probes. This joint modulation of the sensory volume is referred to as acoustic gaze (*sensu* Wisniewska *et al*. (2012)). Echolocating bats in pursuit of an insect represent a dramatic example of acoustic gaze modulation over the span of a few hundred milliseconds - starting out with high, to gradually decreasing source-levels while approaching the insect. Finally, just before interception, call directionality is widened (Jakobsen and Surlykke, 2010; Matsuta *et al*., 2013a). Similar behaviours are seen even in aquatic echolocators like porpoises and dolphins (Ladegaard *et al*., 2017; Wisniewska *et al*., 2012). Studying acoustic gaze and its dynamics provides a direct insight into the sensory decisions of active sensing animals, and is a unique system provided by echolocators.

Studying acoustic gaze and quantifying the underlying acoustic parameters (hence referred to as acoustic gaze inference) is however not a trivial task. The design of microphone or hydrophone arrays in acoustic gaze experiments reflects a multi-variate tradeoff. The ideal (and unrealisable) acoustic gaze inference setup would perhaps have thousands of microphones placed tightly together wherever the animal’s emissions can be captured. Microphone arrays used in real experimental settings however reflect an implicit trade-off between cost, logistics, and current hardware state-of-the-art. Often the limiting factor is the number of recording channels. Given a fixed number of microphones - a tightly spaced array is perhaps able to provide high-angular resolution of the beam aim, but only in a limited volume. On the other hand, intuition seems to suggest that a set of microphones placed wide apart, but covering the overall volume better may perhaps provide a coarser estimate of beam aim - but fail in other parameters. There remains no common framework to understand what kind of array geometry is ideal to perform acoustic gaze inference. Different microphone array geometries may have different abilities to infer the underlying parameters, but the inherent uncertainty in the estimation itself is rarely characterised, for instance through simulations ( but see (Kounitsky *et al*., 2015; Ladegaard *et al*., 2017; Wisniewska *et al*., 2015)). The challenges in acoustic gaze inference is revealed by the kinds of experiments in published studies. Acoustic gaze has been studied in captive settings, where the microphone array and animal movement can be arranged to maximise data collection (e.g. increase the number of emissions recorded by training or constraining animal movement within an arena) (Beedholm *et al*., 2021; Ghose *et al*., 2006; Jakobsen *et al*., 2013; Kloepper *et al*., 2018, 2012; Kounitsky *et al*., 2015; Macaulay *et al*., 2020; Matsuta *et al*., 2013b; Wisniewska *et al*., 2012; Yamada *et al*., 2016; Yovel *et al*., 2010)).Acoustic gaze experiments in the field are equally numerous (Fujioka *et al*., 2016; Jakobsen *et al*., 2015; Ladegaard *et al*., 2017; Motoi *et al*., 2017; Møhl *et al*., 2003; Olsen *et al*., 2020; Surlykke *et al*., 2009). A common theme across both lab and field studies is the potential bias in the kinds of acoustic gaze behaviours that will be uncovered. Echolocators probe their environment differently based on whether they are indoors or outdoors (Surlykke and Moss, 2000), and moreover in lab studies - there is a clear taxonomic bias towards smaller, tamable study species like bats, porpoises and dolphins which can be easily kept in comparison to the larger, wilder counterparts (*e.g.* oilbirds, sperm whales).

Having collected the data, the challenges in performing inference itself become clear when we look at the range of quality criteria applied to the data. Studies perform analysis only on calls/clicks that are aimed within a specific region of the microphone array (Ghose *et al*., 2006; Jakobsen *et al*., 2013; Jakobsen and Surlykke, 2010; Kounitsky *et al*., 2015; Ladegaard *et al*., 2017; Wisniewska *et al*., 2015; Yamada *et al*., 2016). This data curation is done with the understandable goal of only analysing data where parameters can be estimated reliably. This curation step leads to a majority of clicks or calls being discarded, thus resulting in an undersampling of exhibited acoustic gaze. Aside from the undersampling, studying calls/clicks that fall in a specific region while discarding the rest comes with the implicit assumption of uniformity, that the ‘curated’ calls/clicks are the same as those calls/clicks that did not pass the curation. Moreover, current acoustic gaze studies often focus only on the frontal portion of the sound emission - effectively throwing away an entire hemisphere of acoustic data that could potentially be informative too (but see Macaulay *et al*. (2020)). For echolocators, the echoes returning from the frontal half of their emission is indeed the most relevant - however, from a research perspective being able to infer the ‘shape’ of an animal’s acoustic beam all around opens up new venues of research. When the all-round beam-pattern of an animal’s emission is quantified, interesting acoustically mediated behaviours like social interactions or sound propagation in the environment can be studied.

As a parameter-inference problem, acoustic gaze inference as currently performed presents a workflow with ambiguous consequences. Currently acoustic gaze parameters are estimated in a multi-step manner. The sound levels (microphone received levels or unit distance compensated levels e.g. 1m) from spatially sparse mics are interpolated, smoothed or curve-fit in one or two dimensions. The peak of the generated line or surface is then taken to be a higher spatial resolution estimate of the animal’s acoustic beam aim. Once the beam aim is estimated, further click/call parameters such as source-level and beam-width are then estimated (Jakobsen and Surlykke, 2010; Kloepper *et al*., 2018; Ladegaard *et al*., 2017; Wisniewska *et al*., 2015; Yovel *et al*., 2010). I hereby refer to this two-step parameter estimation as the ‘serial’ acoustic gaze pipeline. The serial acoustic gaze pipeline is a set of community-consensus steps that have not been investigated in great detail (but see Guarato *et al*. (2011) and Ladegaard *et al*. (2017)), and represent a heuristic analysis pipelines that researchers have converged upon. Interpolation and curve-fitting of sound levels may perhaps be reasonable when the animal is close to the array or the emission itself has a smoothly varying beam. However, acoustic radiation patterns in general are not guaranteed to smoothly vary, even in theoretical source models (Beranek and Mellow, 2019a). In addition, at least one study (Beedholm *et al*., 2021) has shown interpolation methods themselves can generate false positives, particularly with regards to beam aim. On top of interpolation, another heuristic seen is to discard, or linearly impute microphone received levels when outliers are found (Kounitsky *et al*., 2015; Yovel *et al*., 2010). An alternate to the ‘serial’ pipeline, is to *simultaneously* estimate beam-aim along with source-level and directionality parameters (hence referred to as simultaneous acoustic gaze inference). Guarato and Hallam (2009) began work in simultaneous acoustic gaze, and incorporated beam-aim as a free-parameter. Guarato *et al*. (2011) then compared the performance of the estimated vs. interpolated beam-aim pipelines, and found simultaneous inference recovered ground-truth measurements a majority of the time. Initial simultaneous inference approaches (Guarato and Hallam, 2009; Guarato *et al*., 2011) output only point-estimates, and do not provide the uncertainty of the estimate itself. Additionally, as in any ‘plain’ optimisation based parameter inference, false positives are unavoidable (e.g. unrealistic beam-aim estimates, that also affect all other parameters estimated) due to the lack of a way to incorporate experimental or biological context. Heuristic decisions (whether serial or simultaneous acoustic gaze) likely arise from the reasonable motivation of a good data-model fit. Each of the individual or combination of these heuristics in the analysis pipeline may seem correct from the perspective of researcher intuition, however they can often have systematic effects that can go against the researcher goal at hand (Gelman *et al*., 2026).

In this paper I present a generalised conceptual approach using a Bayesian model-based paradigm. Using a variety of simulated scenarios I show how the Bayesian model-based paradigm overcomes the reliance on data-curation and interpolation/curve-fitting, while also releasing constraints on where the animal must be aiming its beam to necessitate succesful acoustic gaze inference. Finally, I demonstrate the ability to incorporate the arbitrary complexities of real-life scenarios into acoustic gaze inference such as measurement errors in source localisation or received levels. I quantify the utility of the the model-based paradigm with three simulation case-studies outlined briefly below.

### Case study 1. Releasing constraints on acoustic gaze inference

A major data constraint applied to almost all studies of acoustic gaze in experimental settings is that the animal must face, or move towards the microphone array. This is done in order to capture the on-axis emission energy of the call or click. This constraint is reasonable, and also realisable in a context where animals can either be trained/funneled to approach a target (Kloepper *et al*., 2018; Kounitsky *et al*., 2015; Wisniewska *et al*., 2015) or food source (Ghose and Moss, 2003, 2006; Jakobsen and Surlykke, 2010; Ladegaard *et al*., 2017). However, applying such orientation based data filtering means a large set of clicks/calls are discarded. For an ecological and temporally contextual understanding of acoustic gaze, it is important to consider as much of the animal’s sensory behaviour as possible. Especially in the wild, discarding behavioural data that do not fit a spatial constraint can censor a behavioural sequence and induce confounds. For instance, microphones are invariably placed on walls or bulky frames - meaning that any acoustic gaze behaviour studied with current heuristics will automatically only study the incoming flight. The incoming echolocation calls will reveal the properties of acoustic gaze that also carries some form of a clutter response. In contrast, studying acoustic gaze during the incoming and outgoing flight means the ability to capture the natural variation in call/click parameters as a clutter-response emerges, and wanes.

### Case study 2. Post-hoc calibration of acoustic gaze inference beyond the curated range

How well can acoustic gaze be quantified given a microphone geometry, animal location, and measurement noise? This is an open question that always needs to be answered on a case-by-case basis, with various attempts at using proxies like angular resolution based on inter-mic distance from a location (Wisniewska *et al*., 2015; Yamada *et al*., 2016; Yovel *et al*., 2010) or Monte-Carlo simulations (Kounitsky *et al*., 2015; Wisniewska *et al*., 2015). To date, microphone arrays have been planar along the X or Y axis, tightly-spaced (Jakobsen and Surlykke, 2010; Wisniewska *et al*., 2012), in geometries with regular (Ghose *et al*., 2006; Wisniewska *et al*., 2015) or irregular configurations (Fujioka *et al*., 2016; Kounitsky *et al*., 2015). Moreover, the number of channels used vary extensively from 8-48. Beam aim is an important parameter while inferring beam properties, and this has often been separately estimated through various forms of data processing (interpolation, smoothing, curve-fitting or vector-addition) (Ghose and Moss, 2003; Jakobsen and Surlykke, 2010; Kounitsky *et al*., 2015; Wisniewska *et al*., 2015) of microphone received levels or distance compensated levels. The intuition for such interpolation is clear - the beam is strongest on-axis, and when microphones on a plane reveal a convex pattern of received levels, the peak must be the direction in which the call is aimed at. The robustness of this peak-based beam-aim is not clear, extreme-value statistics may be sensitive to noise, while curve-fitting assumes a single-peaked smooth source. In contrast, a Bayesian model-based acoustic gaze inference allows for a careful post-hoc calibration of the total pipeline comprising of the choice of source model, microphone array, animal position, intermediate analysis decisions, and the final outcome parameter uncertainty.

### Case study 3. A priori calibration - maximising acoustic gaze inference abilities while planning data collection

Before any data collection has begun there are many important decisions that an experimenter faces. How many microphones are enough? Where, and how should they be placed? Perhaps most importantly, how well can acoustic gaze be captured given that the animals may fly by at certain distances and orientations? A model-based approach is especially useful as it allows a) estimating acoustic gaze inference ability and b) identifying the conditions under which an array can provide the best parameter inference. Performing an a-priori calibration exercise forces the experimenter to define what sort of parameter uncertainty is acceptable for the scientific goal at hand (*e.g.* 3 dB credible interval for source levels, 10*^◦^* for beam aim). Most importantly however, calibrating the performance of a microphone array under different simulated scenarios allows the experimenter to arrive at the field with an intuition of how to manage the already unpredictable data that is to be collected: where to place the array with reference to the animal/s, and at what distances or orientations they should be recorded at for maximal behavioural sequence continuity.

## II. METHODS

### A. General acoustic gaze inference modelling framework

The general structure of the acoustic gaze inference in the Bayesian framework is as follows:

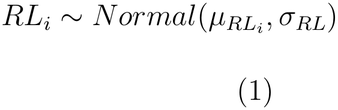

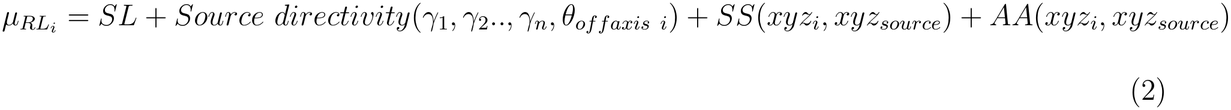

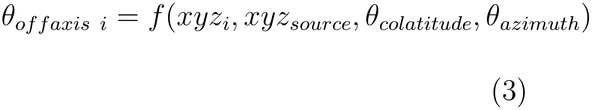

The measured received level *RL_i_* at the *i*th sensor is modelled as the outcome of a Normal distribution whose mean *µ_RLi_* is predicted by the outcome of a combination of the predicted on-axis source level *SL*, the source directivity and spherical spreading *SS*. Unlike previous approaches, it must be stressed here that the input data for this framework is indeed the raw received level (e.g. dB rms, dB SPL, etc.) as measured at the microphones, without any previous distance compensation. The standard deviation *σ_RL_*represents the underlying residual in model prediction due to errors in measurement and data-model mismatch. *θ_offaxis i_* is the relative off-axis angle between the on-axis beam-aim and the microphone position, while *SS* is the spherical spreading (20*log*_10_*radial distance* re. 1m) and *AA* is the atmospheric absorption (*α_absorption_ × radial distance*) expected between the source position *xyz_source_* and the *i*th sensor position located at *xyz_i_*. The chosen source-directivity model may require multiple parameters *γ*_1_*..n* that may be partially known or to be estimated fully.

In this framework, the parameters to be estimated are the on-axis source level *SL*, beam-aim (*θ_colatitude_, θ_azimuth_*) and source-directivity model dependent parameters *γ*_1_*..γ_n_*. Priors for the source level can be formulated for example as truncated Normal distributions based on the available literature (above a minimum expected threshold with decreasing probability for extremely high source-levels). Priors for beam-aim can be physically and biologically constrained to be within a certain range of the movement direction. The measurement standard deviation *σ_RL_*can be estimated from the data itself. The source-directivity model’s parameters *γ*_1_*..γ_n_*may be a mix of known, and to-be estimated variables. Let us take the piston in an infinite baffle as a candidate model. The piston model requires the wave number *k* and piston radius *a* to generate predictions. The wavenumber (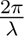, where *λ* is the wavelength) is known *a-priori* as a particular frequency or frequency band is used to quantify received levels. The piston radius *a* however is to be estimated. In other models, e.g. the Gaussian beam model, only the standard deviation of the beam *σ_beam_* needs to be estimated.

The received level *RL* at the sensors, position of the echolocator *xyz_source_* and microphones *xyz_i_* is assumed to be known.

#### 1. Source-directivity models

The formulation of the various source directivity models used here are as follows. The Gaussian model is given by:

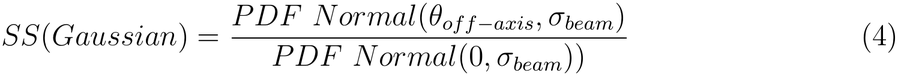

where *σ_beam_* is the standard deviation of the source-beam in decibels. The relative off-axis level is described by a ratio of the on-axis to the off-axis probability density function (PDF). Here, a higher *σ_beam_*leads to a more omnidirectional beam.

The ‘Giuggioli’ model (Giuggioli *et al*., 2015) is a cardiod-type model given by:

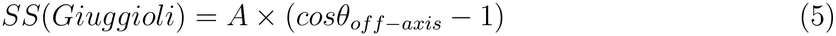

where *A* is the asymmetry parameter. Higher values of *A* correspond to more directional beams.

The piston in an infinite baffle model is a canonical model (Beranek and Mellow, 2019b) whose directivity is described in the following manner:

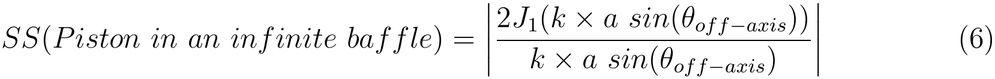

where *J*_1_ is the Bessel’s function of the first kind of order 1, *k* is the wavenumber ( 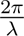, where *λ* is the wavelength), and *a* is the piston radius. The product *ka* predicts the directionality of the resulting beam, with higher *ka* corresponding to more directional beams.

All source-directivity models quantify the off-axis to on-axis ratio, and in the following models that are constructed the 20*log*_10_ of the ratios are used to model the final received levels.

### B. Case study 1. Releasing constraints on acoustic gaze inference

To demonstrate flexible acoustic gaze inference in the field with a sparse microphone array, we simulate an arbitrary array geometry with 20 microphones placed at two heights in an L-shaped configuration. The simulated bat-like echolocator is assumed to emit 4 calls (a number chosen for simplicity of presentation) as it first flies into the ‘L’ shaped array, and turns away from it (Fig. 1A). The echolocator varies its source-level, beam directivity and beam aim on a call-to-call basis, which we try to recover.

**FIG. 1.**
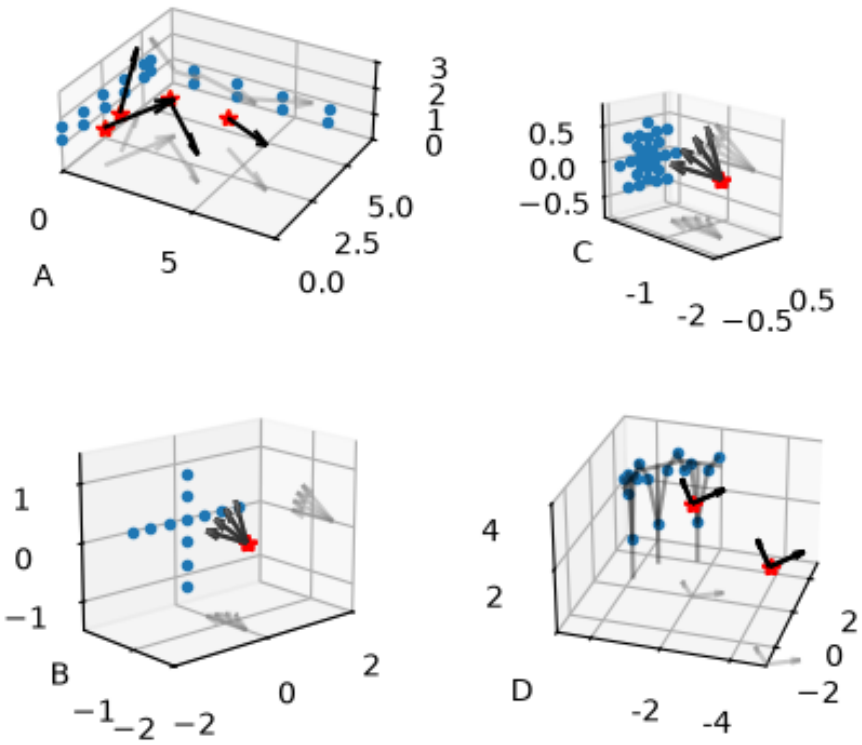
Schematics of the various acoustic gaze inference scenarios simulated. Arrows indicate 3D beam aim, with 2D projections wherever possible, Red stars are echolocator positions, blue dots are microphone/hydrophone positions. A) An echolocator calls into and away from a microphone array in the wild. Four calls are simulated with different colatitudes and elevations that do not align cleanly with the flight direction. Two of the four calls also do not even face the sparse microphone array. B) An echolocator with a medium directivity calling at a variety of off-axis angles onto a 12 channel cross-shaped array C) Echolocator with a high directivity calling at a variety of off-axis angles onto a 48 channel array with a dense square central portion comprising of 25 channels. D) Echolocator with low directivity calling at two different distances (near, far), and two different orientations (facing into, and away) from the array.

The sound directivity was modelled with a Giuggioli-type model. Source-levels were set to initially decrease, followed by an increase at the end. Directivities themselves were set to initially be high, followed by two calls with the same directivity, and one wider beam as the echolocator leaves the vicinity of the reflective array. Atmospheric attenuation was set to 1.5 dB/m. The beams were set to not co-incide with the direction of flight to set a high bar for the current workflow. The parameters of the simulated calls are given in Table 1.

**TABLE I.** Simulated call parameters for the echolocators trajectory shown in Fig. 1. Source level is in dB SPL re 20*µ* Pa. Source directivity is modelled using the Giuggioli model, which requires the asymmetry parameter A. For multi-model comparison, the effective beamwidth (one-sided 6dB drop) in degrees is also provided.

| Call number | Time, s | A | Beamwidth (6dB, deg) | Colatitude, deg. | Azimuth, deg. | Source level, dB SPL | Noise SD, dB | Source X | Source Y | Source Z |
|---|---|---|---|---|---|---|---|---|---|---|
| 1 | 0.0 | 15 | 52 | 90 | 70 | 100 | 1.5 | 1 | 1.0 | 1.45 |
| 2 | 4.5 | 10 | 66 | 70 | 110 | 90 | 1.5 | 1 | 2.0 | 1.50 |
| 3 | 6.0 | 10 | 66 | 100 | -40 | 80 | 1.5 | 2 | 3.5 | 1.60 |
| 4 | 8.0 | 7 | 81 | 80 | -35 | 110 | 1.5 | 4 | 4.0 | 1.20 |

In any kind of real world setting, the underlying source directivity model is not known to the experimenter. Two levels of model ‘ignorance’ were implemented. The first, optimistic scenario is when the experimenter happens to choose a good source-model - mimicked by choosing the Giuggioli model. The second level is when the experimenter chooses a model that only partially recovers beam patterns, mimicked by choosing the Gaussian model. The beam aim and source-level priors for each call *j* are as follows:

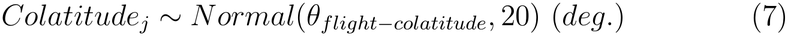

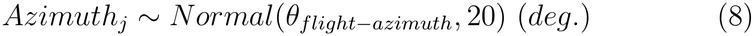

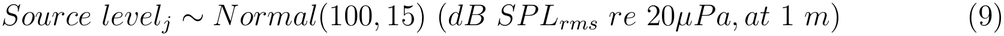

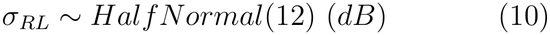

For the Giuggioli model we choose the priors for the asymmetry parameter *A* were set to be *A ∼ Half Normal*(10). The prior for *A* matches the order of magnitude chosen for e.g. *Myotis daubentonii* (Giuggioli *et al*., 2015). For the Gaussian model the beam *σ_beam_* is modelled as *σ_beam j_ ∼ Half Normal*(3) (*dB*). The prior of the Gaussian beam directivity *σ_beam j_* corresponds to a standard deviation of 3, which implies an omni-directional beam - with increasing probability for more directional beams (as expected with bat-like echolocators). The beam’s colatitude and azimuth are centred on the actual flight path direction vector of the nearest interpolated trajectory point and the emission point. To approximate flight direction, the four call emission points were interpolated with a PCHIP shape-preserving interpolator (Virtanen *et al*., 2020) for each dimension independently. The standard deviation of both colatitude and azimuth of the beam was generously set to 20 degrees - indicating a large amount of flexibility in all directions (prior mean *±*40*^◦^*2SD range). Finally, the priors for the call source level was based off commonly reported for bat-like echolocators in the literature - the 2SD extremes of which correspond to ∼70-130 dB SPL at 1m - corresponding to the general range occupied by ‘whispering’ and open-space aerial echolocators (Jakobsen *et al*., 2021). A No-U-Turn sampler (NUTS) was used to run four Markov Chain Monte Carlo (MCMC) chains with 1500 tuning and 500 draws during the sampling. 500 samples were drawn for posterior predictions.

### C. Case study 2. Post-hoc calibration of acoustic gaze inference beyond the curated range

Here I recreate two experimental setups and demonstrate how the Bayesian model-based paradigm can justify the behavioural range over which data may be analysed with two existing array configurations (Jakobsen and Surlykke, 2010; Wisniewska *et al*., 2015). These two experiments were chosen as they represent the two broad ranges of setups. Jakobsen and Surlykke (2010) represents a few-sensor (12 mics) array set up in a relatively spatially sparse manner to study bats with moderate directionality, while Wisniewska *et al*. (2015) represents a high-sensor (48 hydrophones), high-density setup used to study the rather directional porpoise clicks. Both studies apply stringent beam-aim criteria and only analyse calls/clicks that fall within an angular range of the array centre to ensure quality of parameter estimates. Here I investigate the consequences of array-geometry, beam directionality and inference model on the ability to recover ground-truth parameters. In this set of simulations, ‘incorrect’ source models are intentionally used for parameter inference to mimic what happens in real-life scenarios when experimenter ignorance of the true source-model. To validate the long-run reliability of parameter inference I perform parameter recovery tests by repeatedly running inference on multiple simulated datasets. The purpose of parameter recovery checking is to see if the true underlying parameters can be recovered given the current sampling regime (e.g. number of microphones, their arrangements, model choice). In specific I vary the beam-aim parameter to understand how parameter recovery is influenced.

#### 1. Sparse array with intermediate directionality

In Jakobsen and Surlykke (2010), a cross-shaped array of 12 microphones with 0.4 m inter-mic spacing was used to study the acoustic gaze of echolocating bats (*Myotis daubentonii*) as they approach a prey insect. The simulated bat was set to be at 1.5 metres in front of the central microphone (x,y,z: [0,-1.5,0]), emitting a call at 95 dB SPL re 20*µ*Pa (at 1m). Jakobsen and Surlykke (2010) analyse only calls that fall between *±* 10*^◦^* of the centre (corresponding to *±* 80*^◦^* colatitude and azimuth). Four beam-aim scenarios were simulated ranging from on-axis (colatitude,azimuth:[90,90]) to increasingly off-axis (colatitude,azimuth: [80,80],[70,70],[60,60]). Error in received level measurement was simulated by a Normal distribution centred at 0 with 0.5 dB standard deviation. A Gaussian source model with 0.7 dB standard deviation was used, which corresponds to a one-sided -6 dB drop of ∼45 degrees (matching that reported in Jakobsen and Surlykke (2010)). A Gaussian was used instead of a piston model as the piston model is only known to describe the main lobe of a bat call well, with a clear lack of the side-lobes predicted by the piston model (Hartley and Suthers, 1989). Atmospheric attenuation was set to 1.5 dB/m. To mimic the experimenter ignorance of beam geometry in real data, two separate source models were used for parameter inference: the Giuggioli and piston models. 40 independent simulation runs with the same parameter values were generated with unique measurement noise. The priors for the directivity-independent parameters were:

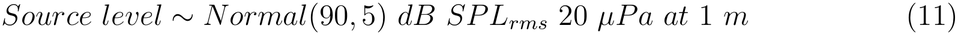

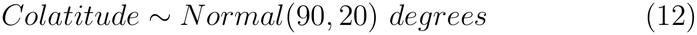

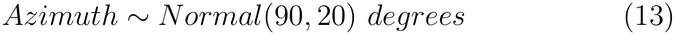

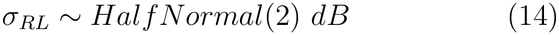

The piston radius prior was *a ∼ Exponential*(1*/*2 *×* 10*^−^*^3^) 1*/m radius*.

The wavenumber *k* = 1016, equivalent to that of the peak frequency 55 kHz. Priors for the source level capture the overall range of source levels reported in the paper. Priors for piston radius *a* capture the order of magnitude of generally reported values, while also encoding an overall lower probability for larger piston radii in the exponential distribution. For the Giuggioli model, the asymmetry parameter *A* was modelled as *A ∼ Exponential*(1*/*6). *A* = 6 was chosen as the rate parameter for the exponential distribution as it matched the 6dB one-sided beamwidth value of ∼90 *^◦^* that a 2 mm radius piston chosen in the other model has. For every simulation run, a No-U-Turn sampler (NUTS) was used to run four Markov Chain Monte Carlo (MCMC) chains with 2000 tuning and 1000 draws during the sampling. 500 posterior prediction samples were also drawn for comparison with the observed received levels.

#### 2. Dense array with directional beams

Wisniewska *et al*. (2015) study the highly directional beams of harbour porpoises (*Phocoena phocoena*) using a series of hydrophone arrays of varying number and geometry. Here we will study the capabilities of the 48 channel ‘small’ array as it represents the best-case scenario for acoustic gaze estimation. The source was set to be 1.5 metres away from the central hydrophone (x,y,z: [0,1.5,0]), at the depth of the central hydrophone (Fig. @ref(fig:wisniewskaresults)A). Four beam-aim scenarios were simulated ranging from on-axis (colatitude,azimuth:[90,90]) to increasingly off-axis (colatitude,azimuth: [80,80],[70,70],[60,60]). A Gaussian beam was simulated with 0.2 dB standard deviation, which corresponds to ∼9*^◦^* (3dB one-sided width) as reported in Wisniewska *et al*. (2015) at a distance of ∼1.5 m. Atmospheric attenuation was set to 0.04 dB/m. A Gaussian source model was used instead of a piston to simulate the source as to date there are no reports of side-lobes as predicted by the piston model (Beedholm and Møhl, 2006; Koblitz *et al*., 2012). Source level was set to 150 dB rms eq. (at 1m re 1*µ*Pa). Measurement noise was simulated by a Normal distribution with 0 mean and 0.5 dB standard deviation.

The priors used were:

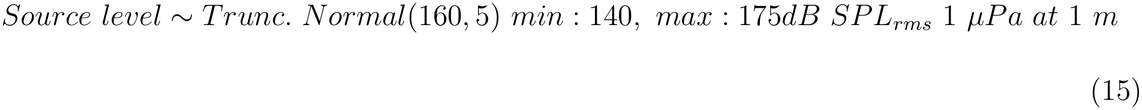

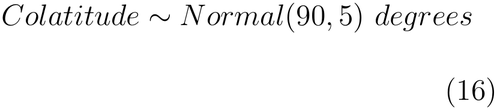

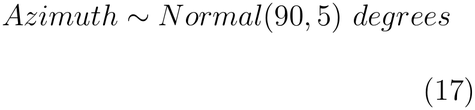

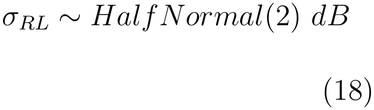

The prior for piston radius was *a ∼ Trunc. Normal*(0.05, 0.03) *min* : 0.001*, max* : 0.083 *×* 3 (*m*) based on the range of piston radius for the least squares error fit in Wisniewska *et al*. (2015). A *k* of ∼544 was used, corresponding to the peak frequency of 130 kHz of clicks reported in (Wisniewska *et al*., 2015). While modelling the Giuggioli model, the asymmetry parameter *A* was modelled as a truncated Normal where *A ∼ Trunc. Normal*(10^3^, 6 *×* 10^3^) *min* : 20*, max* : 20*×*10^3^ to match the beamwidths used for the piston model. The priors for source level was based on Koblitz *et al*. (2012). Colatitude and azimuth prior were set to lay *±* 10 *^◦^* away (2 SD) from the central hydrophone to mimic the potential angles of click emission as the porpoise swims towards the target in front of the array. 40 independent simulation runs with the same parameter values were generated with unique measurement noise. A Hamiltonian Monte-Carlo sampler failed to reach convergence, and showed persistence divergences despite sampler parameter optimisation, and model verification (Betancourt, 2020; Modrák, 2018) (see Results). As an alternate, four chains of a sequential Monte Carlo sampler were run for 4000 draws for both source models.

#### 3. A priori acoustic gaze inference validation and microphone array design

To demonstrate the a-priori use of model-based acoustic gaze inference for array design I study the ‘oilbird’ microphone array that was recently piloted in a field site (Beleyur & Maldonado-Chaparro (Universidad del Rosario, Bogota) ongoing collaboration). The ‘oilbird array’ is a 16 channel array with 13 microphones placed in a zig-zag fashion along three sides of a trapezoid (Fig. 1D). In this particular scenario, the microphone geometry is a given, but we would like to know whether its use can result in scientifically acceptable acoustic gaze inference. More specifically, I investigate the effect of bird distance (near or far) and orientation (flying into, or parallel to the array). Over the course of the project, oilbirds will be acoustically localised by their clicks using time-differences-of-arrival. Acoustically localising a source with time-difference-of-arrivals has an inherent measurement error which is a function of the relative location and distance to the microphone array (de Framond *et al*., 2023; Wahlberg *et al*., 2001).

The source level of the oilbird was set to 100 dB SPL re 20 *µ*Pa at 1m (Brinkløv *et al*., 2017). Two source-array distances were simulated, the ‘near’ (∼2.5m) and ‘far’ (∼6.5m) from a central microphone. Beam directionality was modelled as a Giuggioli beam, with moderate directionality of *A* = 5, corresponding to a 10 dB smooth front-back drop. Two beam directions were simulated with the oilbird either pointing towards the array (azimuth=35*^◦^*), or parallel to the array (azimuth=-60 *^◦^*). The birds often emerge from the cave and fly up in this field site. To mimic the naturalistic flight behaviour of the birds, the colatitude was set to point slightly upwards (colatitude=70 *^◦^*). Measurement noise was modelled as a Normal distribution with 0 mean and 1 dB standard deviation on each microphone. Localisation error was simulated proportional to distance with 0.1 m (near source) and 0.3 m (far) standard deviation for the XYZ axes independently. For instance, an xyz standard deviation of 10 centimetres each corresponds to a 1 SD euclidean distance of ∼0.17 m, and a 2SD euclidean distance of ∼0.5 m from the original source position. A total of four scenarios were simulated (2 distances x 2 orientation) with 10 simulation runs each. Given the non-analytical nature of the localisation error function, approximate Bayesian computing was used to perform acoustic gaze inference. The distance between simulated and observed data were calculated with a Gaussian distance function 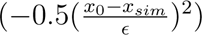, where *ɛ* = 1 for our inference runs, until 1500 draws were obtained.

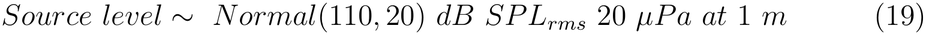

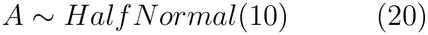

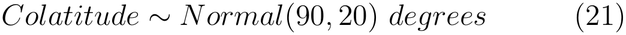

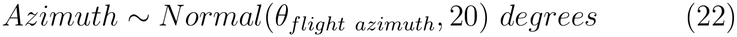

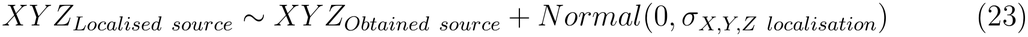

Source level priors match those reported in literature (Brinkløv *et al*., 2017). Given the current lack of click directivity data in oilbirds, a wide prior was used. Colatitude priors actively ignore the known upward direction of flight. The prior mean of azimuthal orientation *θ_flight azimuth_* was set to 45 *^◦^* when the animal was simulated to be flying towards the array, and -45 *^◦^* when flying parallel to the array. *XY Z_Localised source_* was the estimated source position after considering the errors in acoustic localisation, where the overall standard deviation *σ_X,Y,Z localisation_* was known to be either 0.1 or 0.3 m.

All computations were done on Python 3.12.2 (Van Rossum and Drake Jr, 1995), with the PyMC (Abril-Pla *et al*., 2023), NUTPIE No-U-Turn sampler (Seyboldt *et al*., 2026), and arviz inference packages (Kumar *et al*., 2019). When divergences were observed, the step-size was gradually decreased till no more divergences appeared. Chain convergence was verified by low Rhat values less than 1.1. Effective sample sizes for all accepted simulations were set to be at least 5*×N_chains_* when the MCSE values were of acceptable range (Gelman *et al*., 2025) (especially for some of the sequential Monte-Carlo runs, all other inference runs had *≥* 200 effective sample sizes). Prior predictive checks were performed to ensure predictions were in a valid range before posterior estimation. Posterior predictive distributions were compared with observed data to assess model fit, and trace-plots were visualised to ensure even mixing of chains.

## III. RESULTS

The long-run reliability of the inference procedure reported here was estimated by the coverage frequency of the 89% credible highest-density intervals of the posterior distributions of each parameter.

### A. Flexible acoustic gaze inference under field-like settings

All acoustic gaze parameters could be successfully inferred (indicated by the credible intervals covering the ground-truth) in the ‘flexible’ data collection scenario (See Fig.2 A-C for a graphical overview). Even when the data generating model was different from the one used to infer acoustic gaze (as is often the case in any real-world setting) - source level and azimuthal beam-aim estimates matched the ground-truth very closely (Fig.2 D,H). As expected from the limited spatial spread in the Z axis of the microphone array - colatitude estimates (Fig.2 F,J) were much wider than azimuthal estimates (Fig.2E,I). When the in-optimal Gaussian model was used as a source-model, beam-width estimates qualitatively matched (Fig.2K) the ground-truth, but did not accurately recover the temporal trends in call directionality. Overall, beam-width seemed to be the most sensitive to model-choice.

**FIG. 2.**
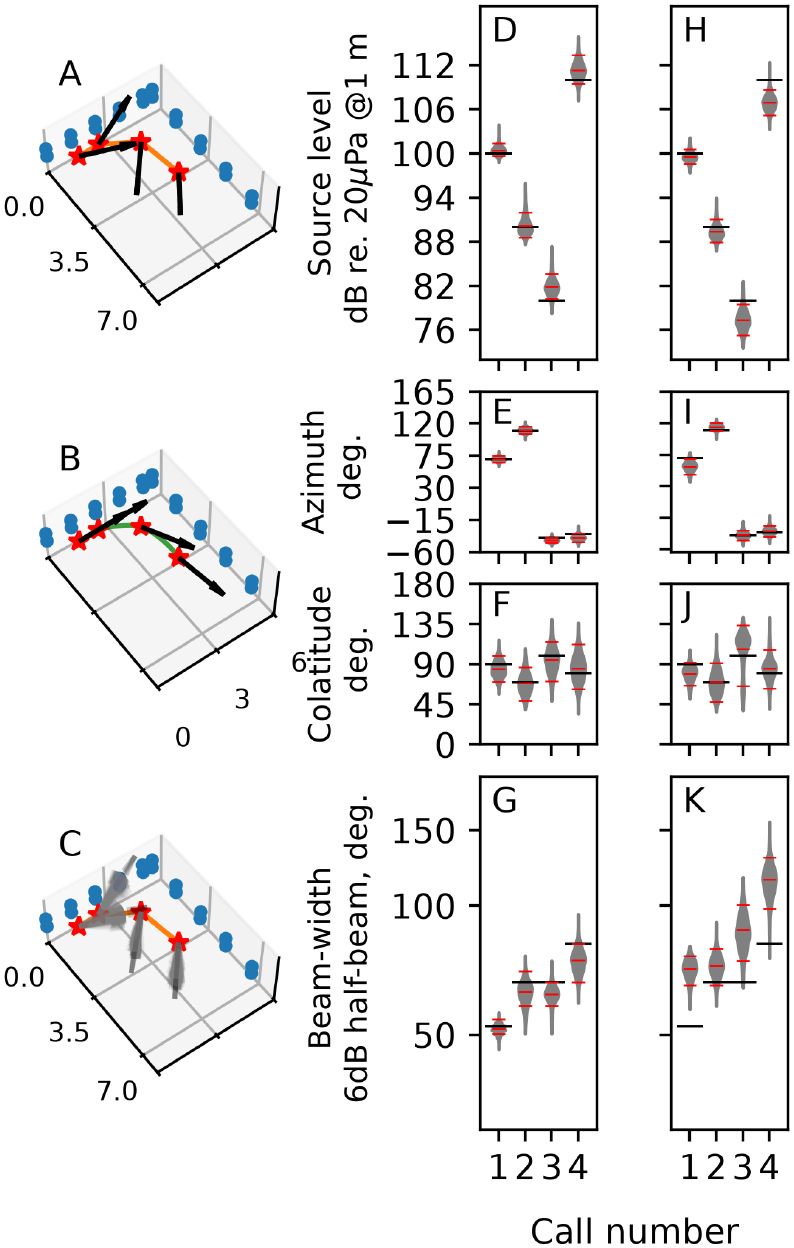
A-C: Data and pipeline for flexible acoustic gaze inference. A) Ground-truth simulation data showing four emission points (calls 1-4 from left to right, red stars), beam-aim (black arrows), and the linearly-connected flight path (orange-lines). B) The priors used for beam-aim (black-arrows) are centred on the interpolated flight path (green line) between emission points. C) Recovery of beam-aim. The posterior distribution for beam-aim with the Giuggioli model (faded grey arrows), and the mean (longer dark grey arrow) show the original beam-aims in A can be recovered. D-G: Posterior distributions when a good source-model (Giuggioli) is chosen H-K: posterior distributions when a mediocre source-model (Gaussian) is chosen. Horizontal black-lines indicate ground-truth values, red lines indicate median and 89% highest-density intervals of the posteriors. G,K have a logarithmic y-axis. Source-level and beam aim can be reasonably recovered independent of source-model. Beam-width quantification requires good model-data fitting.

### B. Post-hoc calibration of acoustic gaze inference beyond the curated range

#### 1. Post-hoc calibration of a sparse array with intermediate directionality

Both Giuggioli and piston models provided satisfactory parameter recovery over a wide range of beam-aims (Fig.3 A-H & Table 2). All parameters except for beam-width showed suffiicient coverage (*∼* 89%) until at least 20 degrees off-axis (colat,azim: 70,70). It is important to keep in mind that strictly speaking we cannot actually expect *≥* 89% coverage given that the data-generating models used for inference (Giuggioli & piston) and the underlying ground-truth models (Gaussian) are actually different. If anything, the ability to reach *∼* 89% coverage is evidence that despite model-mismatch we can recover underlying parameters reasonably well. The piston model showed heavy under-coverage at the most extreme beamaim (colat,azim: 60,60) for all parameters except source-level, while also already showing under-coverage of beamwidth at 20 degrees off-axis. Inference with the Giuggioli model however showed a more reliable coverage of all parameters until 30 degrees off-axis - where it began to falter. Jakobsen and Surlykke (2010) limit their analysis of calls in their analysis pipeline with the piston model to 10 deg. off-centre. Across all beam-aims, source-level uncertainty remained *≤* 2 dB, and beam-aim itself was ∼ *±* 10 *^◦^* for most simulated scenarios. Beamwidth uncertainty was ∼ 7 *^◦^*, while not negligible, still represents a relatively small fraction of the actual beamwidth of 45 *^◦^* (∼15 %). Even though Jakobsen and Surlykke (2010) use a separate analysis pipeline with the piston source model, the model-based inference here provides independent support of the curated angular ranges used by them. For further inspection of parameter recovery results, see Supplementary Information (SI) Fig. S1.

**FIG. 3.**
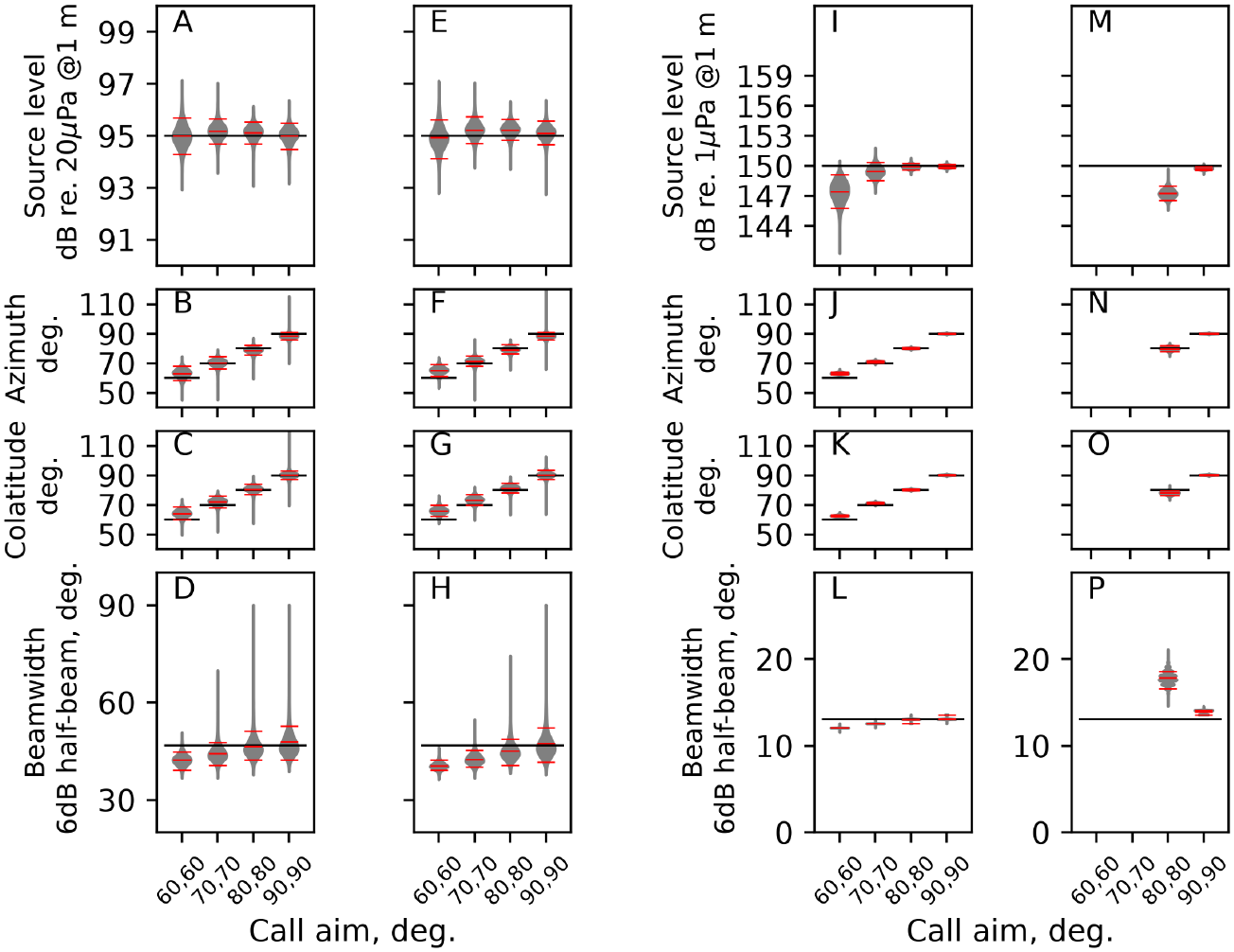
Results of acoustic gaze inference from one simulation run across two different scenarios 1) sparse 12 channel array with medium directivity source simulating a bat call (A-H) and 2) dense 48 channel array with highly directional source simulating a porpoise click. Posterior distributions (grey), means and 89% highest density credible intervals (red lines), and ground-truth (black lines) are shown. Two source models were used for parameter estimation: the Giuggioli (bat simulation:A-D, porpoise simulation: I-L), and piston in an infinite baffle (bat simulation: E-H, porpoise simulation: M-P). In the bat simulation (A-H), both models perform similarly across the range of beam-aims, with the Giuggioli model showing wider coverage across beam-aims. In the porpoise simulation (I-P), the Giuggioli out-performs the piston model in parameter inference. The piston model fails to converge at off-axis beam-aims.

**TABLE II.** Simulated bat parameter recovery results when two mis-specified source models are used to recover acoustic gaze over a series of off-axis beam directions. The Giuggioli model provides reasonable coverage for all parameters till *∼* 20 degrees off-axis, with the piston model only doing so till *∼*10 degrees off-axis. Source-level can be robustly inferred over all beam-aims and both models despite mis-specification. The credible interval coverage for each parameter, and 89% highest-density interval width is reported over 40 simulations for each aim.

| Inf. model | Aim:Azim/Colat | Source level, coverage | Crl range, dB | Beam-width coverage | Crl range, deg. | Azimuth coverage | Azim. Crl range | Colatitude coverage | Colat. Crl range |
|---|---|---|---|---|---|---|---|---|---|
| Giuggioli | 60.00 | 0.97 | 1.45 | 0.38 | 5.34 | 0.70 | 9.97 | 0.80 | 8.81 |
| Giuggioli | 70.00 | 0.97 | 1.00 | 0.85 | 7.42 | 0.95 | 8.56 | 0.88 | 8.51 |
| Giuggioli | 80.00 | 0.92 | 0.90 | 0.97 | 11.12 | 0.95 | 7.31 | 0.90 | 8.19 |
| Giuggioli | 90.00 | 0.92 | 1.10 | 0.95 | 17.21 | 0.95 | 7.18 | 0.97 | 8.62 |
| Piston | 60.00 | 0.97 | 1.44 | 0.00 | 3.04 | 0.35 | 7.84 | 0.38 | 7.71 |
| Piston | 70.00 | 0.90 | 0.99 | 0.45 | 5.43 | 0.85 | 7.25 | 0.82 | 7.69 |
| Piston | 80.00 | 0.95 | 0.86 | 0.97 | 10.47 | 0.95 | 6.54 | 0.90 | 7.52 |
| Piston | 90.00 | 0.95 | 1.04 | 0.90 | 18.15 | 0.90 | 6.70 | 0.95 | 7.76 |

#### 2. Post-hoc calibration of a dense array with directional beams

The original models run with a Hamiltonian Monte-Carlo sampler showed repeated divergences across multiple seeds. Divergences are caused by model misspecification or multi-modality in the data-model combination (Betancourt, 2020; Gelman *et al*., 2026; Modrák, 2018). Coding errors were ruled out by setting tight priors centred around the known value and ground-truth were recovered. The persistent divergences pointed to the weakly-identifiable nature for this particular experimental setup (Betancourt, 2020).

The Giuggioli source model provided reliable parameter inference, in contrast to the piston model which failed to converge on any off-axis simulation data (Fig.3 I-P, Table 3). The Giuggioli model showed appropriate credible interval coverage for all up till 10 degrees off-axis (colat,azim: 80,80). For inference beyond 10 degrees off-axis, while there is undercoverage, the extent of parameter bias is still surprisingly reasonable. Source-level is the main parameter, being under-estimated by 3-4 dB at most - which may be acceptable. In contrast, the piston model shows strong bias already at 10 degrees off-axis, with beam-width over-estimates by ∼4 degrees along with source-level bias of 3-4 dB. Moveover, even in the on-axis scenario, the piston model shows a small but clear positive bias in beam-width. The results here provide independent support for the data curation measures undertaken in Wisniewska *et al*. (2015), who use a piston model in their analysis pipeline, and only analysis clicks that fall ∼2-3 degrees off-centre in the small array. While Wisniewska *et al*. (2015) use a different analysis pipeline, the results here show that the piston model may provide reasonable estimates when used for clicks that are pointing very close to the centre. For further inspection of parameter recovery results, see SI Fig. S2 for all inference runs.

**TABLE III.** Simulated porpoise parameter recovery results when two mis-specified source models are used to recover acoustic gaze over a series of off-axis beam directions. The Giuggioli model provides reasonable coverage for all parameters till 10 degrees off-axis. The piston model only works in a limited manner in the on-axis aim, showing coverage of beam-aim, with small biases in source-level and beam-width estimates. The credible interval coverage for each parameter, and 89% highest-density interval width is reported over 40 simulations for each aim.

| Inf. model | Aim:Azim/Colat | Source level, coverage | Crl range, dB | Beam-width coverage | Crl range, deg. | Azimuth coverage | Azim. Crl range | Colatitude coverage | Colat. Crl range |
|---|---|---|---|---|---|---|---|---|---|
| Giuggioli | 60.0 | 0.1 | 3.0 | 0.0 | 0.0 | 0.0 | 1.7 | 0.0 | 1.3 |
| Giuggioli | 70.0 | 1.0 | 1.6 | 0.4 | 0.2 | 0.4 | 1.3 | 0.1 | 1.1 |
| Giuggioli | 80.0 | 0.8 | 0.5 | 1.0 | 0.2 | 0.9 | 0.8 | 1.0 | 0.7 |
| Giuggioli | 90.0 | 0.9 | 0.3 | 1.0 | 0.4 | 0.9 | 0.5 | 1.0 | 0.5 |
| Piston | 60.0 | 0.0 | 1.9 | 0.0 | 0.2 | 0.0 | 8.0 | 0.0 | 6.9 |
| Piston | 70.0 | 0.0 | 3.5 | 0.0 | 0.0 | 0.0 | 1.5 | 0.0 | 1.7 |
| Piston | 80.0 | 0.0 | 1.4 | 0.0 | 2.2 | 1.0 | 3.7 | 0.2 | 3.5 |
| Piston | 90.0 | 0.3 | 0.3 | 0.0 | 0.4 | 1.0 | 0.5 | 1.0 | 0.5 |

### C. A priori calibration of the oilbird array

The proposed oilbird rig may be able to recover source-level reliably ((Fig.4 A-D), Table 4), followed by beam-aim. The current setup and virtual oilbird locations simulated would fail at recovering beam width. Despite the asymmetric distance-wise localisation error implemented (nearby sources have less localisation error than far-away sources), it is encouraging to see that source-level can be estimated well (∼ 2-5 dB credible interval widths) across the four simulated scenarios. Beam-aim could be reasonably estimated to within the correct quadrant (∼60-100 deg. credible interval widths) - but this uncertainty naturally constrains the spatial resolution of the acoustic gaze inference. Beamwidth estimation however provided extremely wide credible intervals when the bird was far, and extremely biased intervals when near. Neither situation in beamwidth estimation provide any gain in scientific insight (see Fig. S3 for all inference runs).

**FIG. 4.**
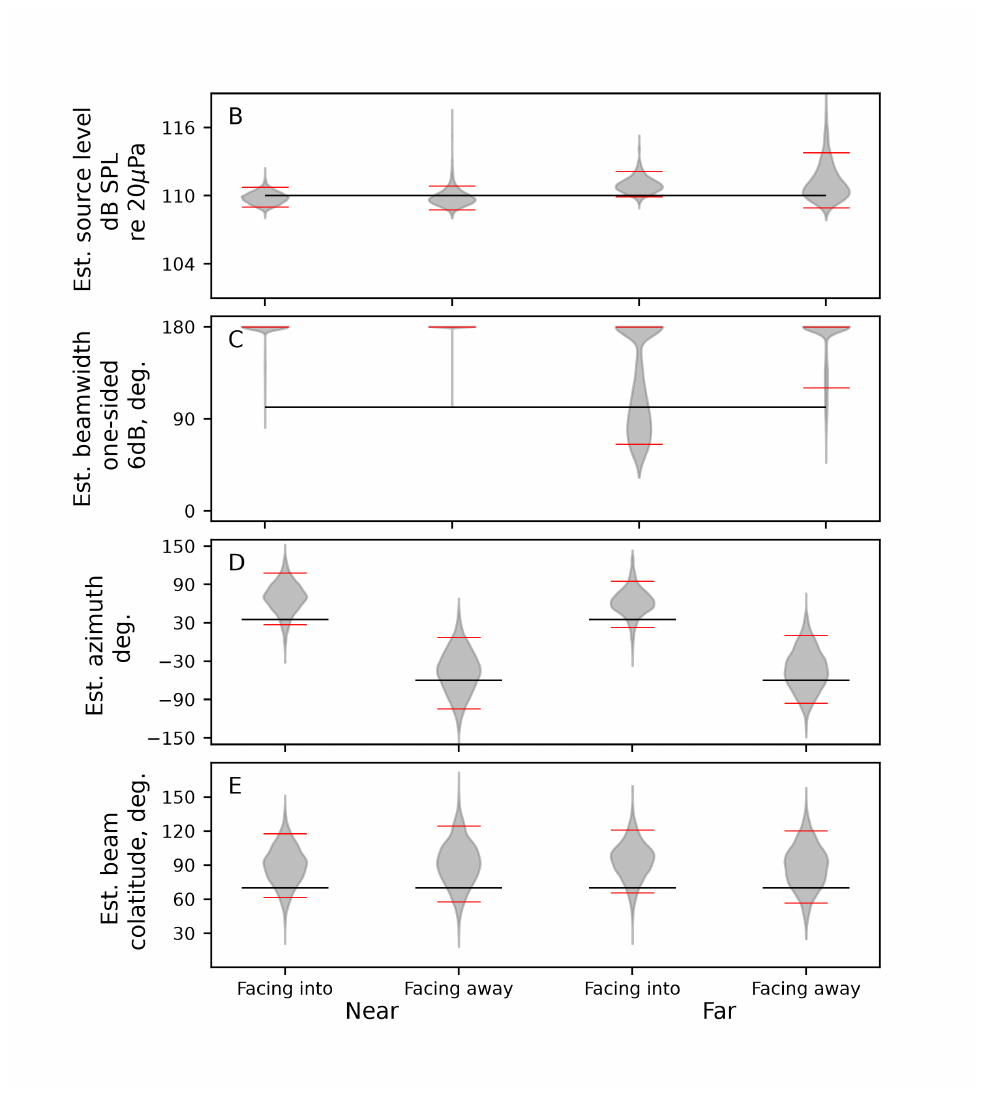
Credible interval coverage of the for a Giuggioli-type omnidirectional source at various distances and orientations to the “oilbird array”. Beam aim and source-level can be recovered even when the source is not pointing towards the array, but beam-width cannot be estimated reasonably at all.

**TABLE IV.** Simulation coverage and mean credible interval widths for beam parameters in the oilbird simulation. Aside from beam-width, most parameters can be recovered reliably most of the time. Azimuth can be recovered reliably, but coarsely to the correct quadrant, while colatitude can be recovered with slightly better precision. A slight under-coverage in source-level is observed when the source is far and pointing towards, but presents a minor biased estimate (See SI).

| Distance | Beam aim | Source level, coverage | CrI range, dB | Beam-width coverage | CrI range, deg. | Azimuth coverage | Azim. CrI range | Colatitude coverage | Colat. CrI range |
|---|---|---|---|---|---|---|---|---|---|
| Far | Parallel | 1.00 | 4.85 | 0.00 | 55.96 | 1.00 | 109.04 | 1.00 | 62.75 |
| Far | Towards | 0.70 | 2.00 | 0.00 | 108.30 | 1.00 | 74.52 | 1.00 | 56.38 |
| Near | Parallel | 1.00 | 2.73 | 0.00 | 0.00 | 1.00 | 101.37 | 0.90 | 61.02 |
| Near | Towards | 0.80 | 1.66 | 0.00 | 33.44 | 1.00 | 73.75 | 1.00 | 57.21 |

## IV. DISCUSSION

In this work I have demonstrated the acoustic gaze inference framework for three purposes: 1) studying acoustic gaze in a behaviourally contextual manner, 2) extending, and *post-hoc* calibration of previously published acoustic gaze studies and 3) *a-priori* systematic planning of acoustic array usage in the field to generate acoustic gaze data with acceptable resolution.

The major determinant of a study is largely not the hands of the experimenter - but rather on the uncontrollable behaviour of the animal’s movement and nature of emitted sound. The model of choice used to perform acoustic gaze inference is however under our influence. Most parameters (source-level and beam-aim) seemed unexpectedly robust to model-choice. The model-based acoustic gaze inference here can recover underlying beam dynamics when emitters are pointing off-axis, and are at arbitrary distances from the microphone array. Moreover, using a multi-model comparison I also show the ability to honestly assess how well the different parameters of acoustic gaze can be recovered. We can now truly study acoustic gaze dynamics in the wild in a quantitative manner, having removed many of the typical ‘curation’ measures (e.g. peak of the beam must fall within *±* X degrees of the centre, animal must be within a certain distance of the array). The biggest advantage in taking the model-based approach is that a sequential analysis of acoustic gaze behaviour can be performed without the need for marginal averaging over calls emitted from similar locations. A general issue with existing acoustic gaze pipelines is the lack of biological/physical constraints into the parameter estimation - leading to occasional ‘outlier’ estimates. For instance in Guarato *et al*. (2011), a few calls seem to be pointed directly at the ceiling - likely resulting from the unconstrained beam-aim estimation used. Unconstrained parameter estimation has the same problems as using non-informative priors in the Bayesian approach - often leading to odd outcomes (Gelman *et al*., 2026).

From the set of four case-studies, one can qualitatively say that beam-width is amongst the harder acoustic gaze parameters to infer. Recovering beam-width seems to require a certain factor of luck in the juxtaposition of source-location, model of choice, beam-aim and mic geometry (*e.g.* see Fig. 2 G,K, where even with the correct model the intervals almost do not cover the groundtruth beamwidth). The exact statistical coverage of the model may not be as interesting to the bio-acoustician - though it indicates which variables can be reliably estimated. Even when a model provides very low coverage, the extent of bias is important to note as it may affect our ability to draw valid conclusions from the observed behaviours. For instance, in the highly directional porpoise simulation (Table 4) there is a low coverage of beamwidth at even 30 degrees off-centre (Fig. 3 I-L), but the extent of bias is surprisingly minimal for all acoustic gaze parameters. Off-centre clicks can thus be interpreted having considered these caveats.

Post-hoc simulation based validation of two acoustic gaze studies showed that experiment-specific investigation is necessary to validate parameter inference. Here I quantitatively demonstrate the abilities of the simultaneous acoustic gaze pipeline in extending the sampling area of published setups. Multiple datasets are often collected with ‘stable’ microphone array systems over time (‘tristar’ arrays (de Framond *et al*., 2023), cross-shaped arrays (Jakobsen *et al*., 2013; Jakobsen and Surlykke, 2010), or room-affixed geometries (Eveland *et al*., 2026; Ghose *et al*., 2006; Ghose and Moss, 2006)). A post-hoc simulation study may be particularly worthwhile to study previously discarded calls/clicks, allowing the study of a wider range of biologically relevant calls/clicks. Even in ‘controlled’ studies where animal’s movements are predictable, increasing the angular range within which calls/clicks are analysed represents a ∼4X increase in the effective area over which inference can be performed for the simulated setups of Jakobsen and Surlykke (2010) and a ∼11X increase for the Wisniewska *et al*. (2015) setup (see Supp. Info for derivation).

The ‘oilbird’ example presented here shows the steps involved in assessing if an experimental setup comprising of microphone array and typically observed behaviours will yield useful scientific insights. When designing a microphone array system from scratch, a priori design allows iterations over multiple array-geometries, until a reasonably well performing version is found. Even without further array-design iterations we now know that the 16 channel ‘oilbird’ array works best when the animal is flying towards the array, and is relatively close. The 16-channel array as it stands cannot provide high-resolution insights into beam-aim and should not be used to infer anything about beam-width. For source-level dynamics however, the array is appropriate. For the larger goals of the oilbird acoustic gaze project however, a new round of iterative array-design is clearly needed. The oilbird use-case highlights the ‘model-first’ approach that pushes the experimenter to imagine the kind of behaviour animals may show, lay out data collection constraints, and finally state the acceptable range of parameter uncertainty for scientific inference. Examples of analogous tools in the world of stereo-camera tracking include the ‘easyCamera’ package (Theriault *et al*., 2014) - which quantifies the 3D tracking accuracy of animals given the orientations and placement of the multi-camera array system. The power of the Bayesian model-based acoustic gaze inference lies in its wide applicability, and generalisability to non-standard conditions. The presented modelling framework can be used in situations where some received level measurements are censored (e.g. noisy/clipped microphones across different calls), or even when there is non-negligible uncertainty in microphone and source positioning. Most importantly, arbitrarily complex source-directivity models can be incorporated - that go beyond the current set of popular directivity models used.

Given the reasonable body of literature on acoustic gaze, the currently proposed approach may seem redundant, especially since the demonstrations are simulated examples. The purpose of keeping this work *in-silico* are two-fold: 1) cleanly demonstrating the ability of this framework to handle complex scenarios that will be encountered in experimental settings and 2) highlighting that the need for systematic explorations of ‘parameter space’ to identify the circumstances under which inference can be performed. Essentially, the ability to obtain acoustic gaze parameters from data do not automatically promise their long-term accuracy or validity. The simulations thus directly demonstrate the need for a principled workflow rather a set of heuristics that recover underlying ground-truth measurements or researcher intuitions (Gelman *et al*., 2026; Wolkovich *et al*., 2026). Only once the computational validity of the procedure have been established can the results from experimental data be reasonably assessed.

In this study, I used analytically tractable source-directivity models to aid fast parameter estimation and chain convergence. In its most general form model-based acoustic gaze inference is not limited to such simple 1-2 parameter models, and can be extended to arbitrary source models too. More detailed analytical models such as a piston in a sphere, cap of a sphere (Beranek and Mellow, 2019a) are alternate options. More complex bi-lobed directivity models have been proposed for bat echolocation calls (Ghose *et al*., 2007), and another option is the rectangular cap of a sphere (Beranek and Mellow, 2019a) that incorporate ventral or dorsal asymmetry in emitted beams, as observed in both bats and whales (Ghose *et al*., 2007; Koblitz *et al*., 2012). Optimised computational implementations for the directivity of some detailed analytical models exist (Beleyur, 2022) - however their use in parameter estimation is hindered by the lack of closed-form solutions to models. Predictions need to be generated through time and computationally intensive numerical integration. Despite optimised code implementations, one parameter set can take anywhere between 5-10 minutes for some of these models. One possible work-around here is to train Gaussian process models on a subset of intensive computational runs (Stetzler *et al*., 2022), and then use the trained Gaussian processes as emulators for acoustic gaze inference.

The presented acoustic gaze inference framework demonstrates the power of releasing assumptions and constraints on data, and taking a ‘model-first’ approach to allow a wider study of active sensing under natural conditions. While the focus of this paper has been on inferring acoustic gaze of echolocating animals, the approach is equally valid to infer the overall beam dynamics of any sound source, biotic or not. Even in the biotic realm, the presented model-based framework could be used to expand the acoustic dynamics of important communication signals like bird song and calls, where intriguing behaviours like opponent-facing and directionality modulation have been observed (Brumm and Todt, 2003; Yorzinski and Patricelli, 2010). Future work could focus on validating the performance of this framework under experimental settings, followed by applying the study of acoustic gaze with arbitrary microphone geometries in the wild.

## V. AUTHOR DECLARATIONS AND DATA/CODE AVAILABILITY

This author declares no conflicts of interest. No generative AI was used in the preparation of this manuscript or associated code. All code and data in this paper are accessed at the following DOI: 10.5281/zenodo.23220591 under a CC BY 4.0 License.

## ACKNOWLEDGMENTS

TB would like to thank members of the Active Sensing Collectives Lab, Felix Häfele, Danuta Wisniewska for discussions, and especially Lasse Jakobsen for detailed discussions and comments. Computations were performed on the HPC systems Raven and Viper at the Max Planck Computing and Data Facility. This work was funded by the Carl-Zeiss Stiftung Nexus Junior Research Group for Active Sensing Collectives (P2023-09-006).

## VI. SUPPLEMENTARY INFORMATION

### A. Increased effective sampling area calculation

The *±* X degrees off-centre limit applied in many studies creates a fictional square within which all calls/clicks have to fall. If we make the assumption that the number of studied calls is proportional to the ‘monitoring’ area we will perform parameter estimation on - what is the advantage provided by the increased angular boundaries from *±* 10 *^◦^* to *e.g. ±* 25 *^◦^* (for the Jakobsen and Surlykke (2010) simulation) and *±* 5 *^◦^* to *e.g. ±* 15 *^◦^* (for the Wisniewska *et al*. (2015) simulation).

The side-length of the fictional bounding-square is given by 2 *× R_bat−centre_tan*(*θ_max_*), where *R_bat−centre_* is the radial distance between the bat and the centre of the array, and *θ_max_*is the angular bounds set by the experimenter. The effective sampling area is thus decided by the distance to the array-centre and the distance of the bat, and is given by:

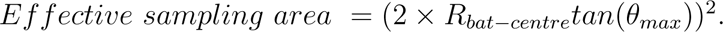

The relative increase in effective sampling area is independent of the distance to the array and is a function of *θ_max_* (proportional to the *tan*(*θ_max_*)^2^ terms), thus the effective sampling area increase is: 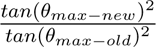. When we substitute the limits used by Jakobsen and Surlykke (2010) (*±* 10 *^◦^*) and this paper (*±* 20 *^◦^*) - we obtain - corresponding to an effective sampling increase of ∼4.

Wisniewska *et al*. (2015) analyse clicks that are within *±* 6-8cm from the centre of the setup for the small array. 6-8 cm from the centre corresponds to 2-3 degrees off-axis. We see that moving from *±* 3 *^◦^* to *±* 10 *^◦^* provides an ∼11 factor increase in effective sampling area.

---

**FIG. S1.**
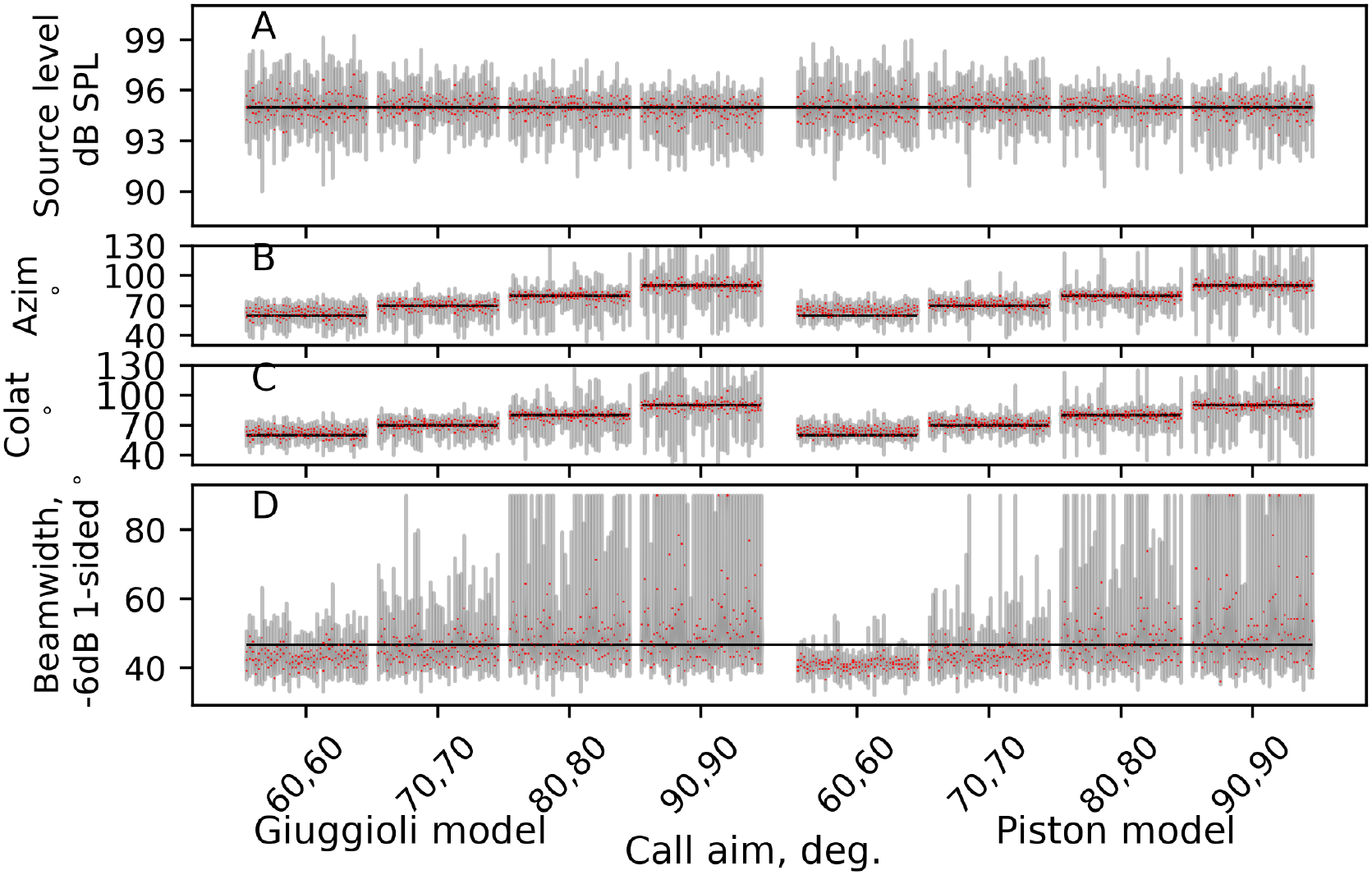
Credible interval coverage of the Giuggioli (left) vs. Piston models (right) for various parameters in a simulated bat call. Both models perform almost equally well, with the Giuggioli showing a generally better performance in capturing the underlying ground-truth.

**FIG. S2.**
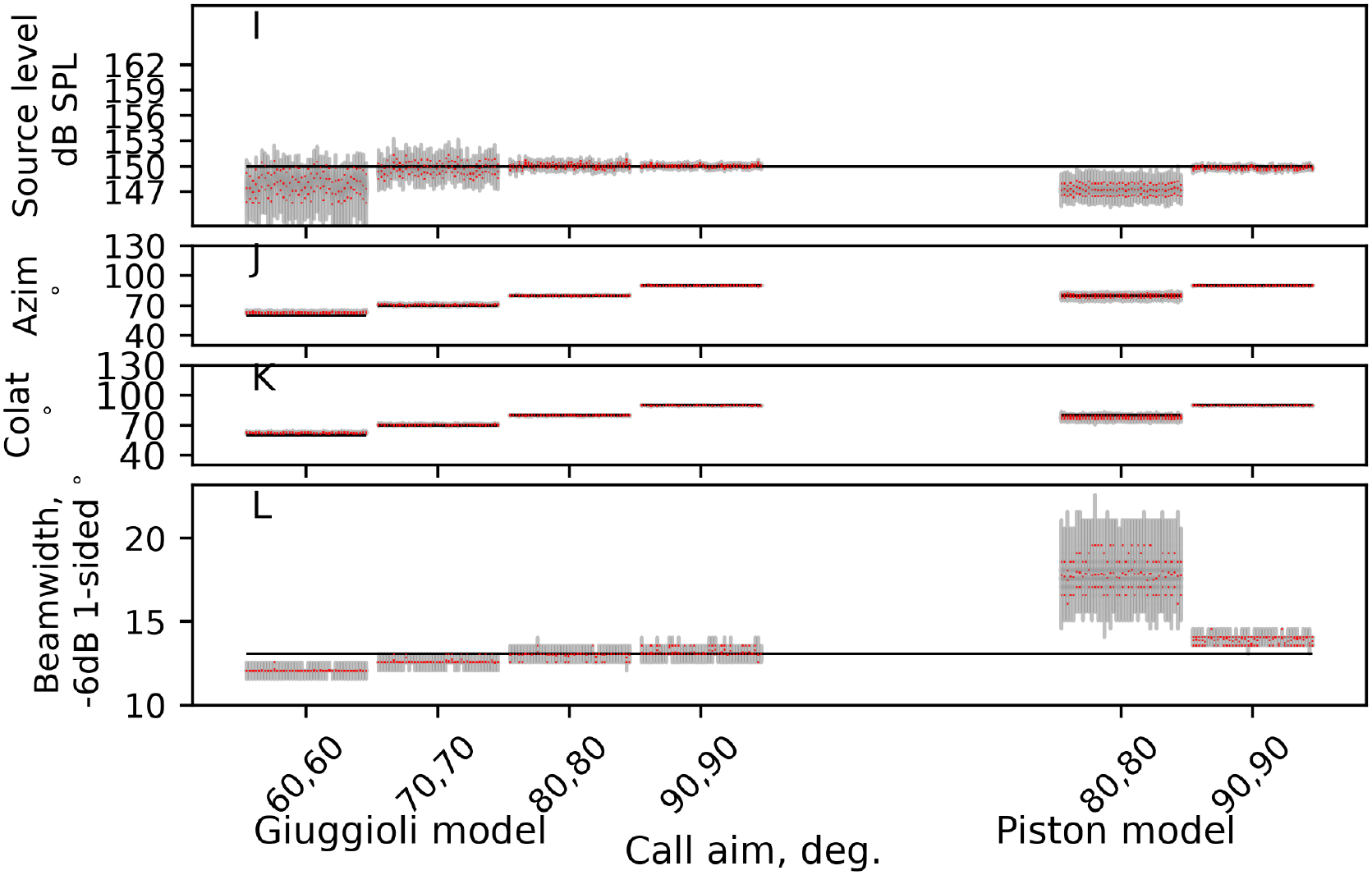
Credible interval coverage of the Giuggioli (left) vs. Piston models (right) for various parameters in a simulated porpoise click. The Giuggioli source-model is superior in capturing the underlying ground-truth.

**FIG. S3.**
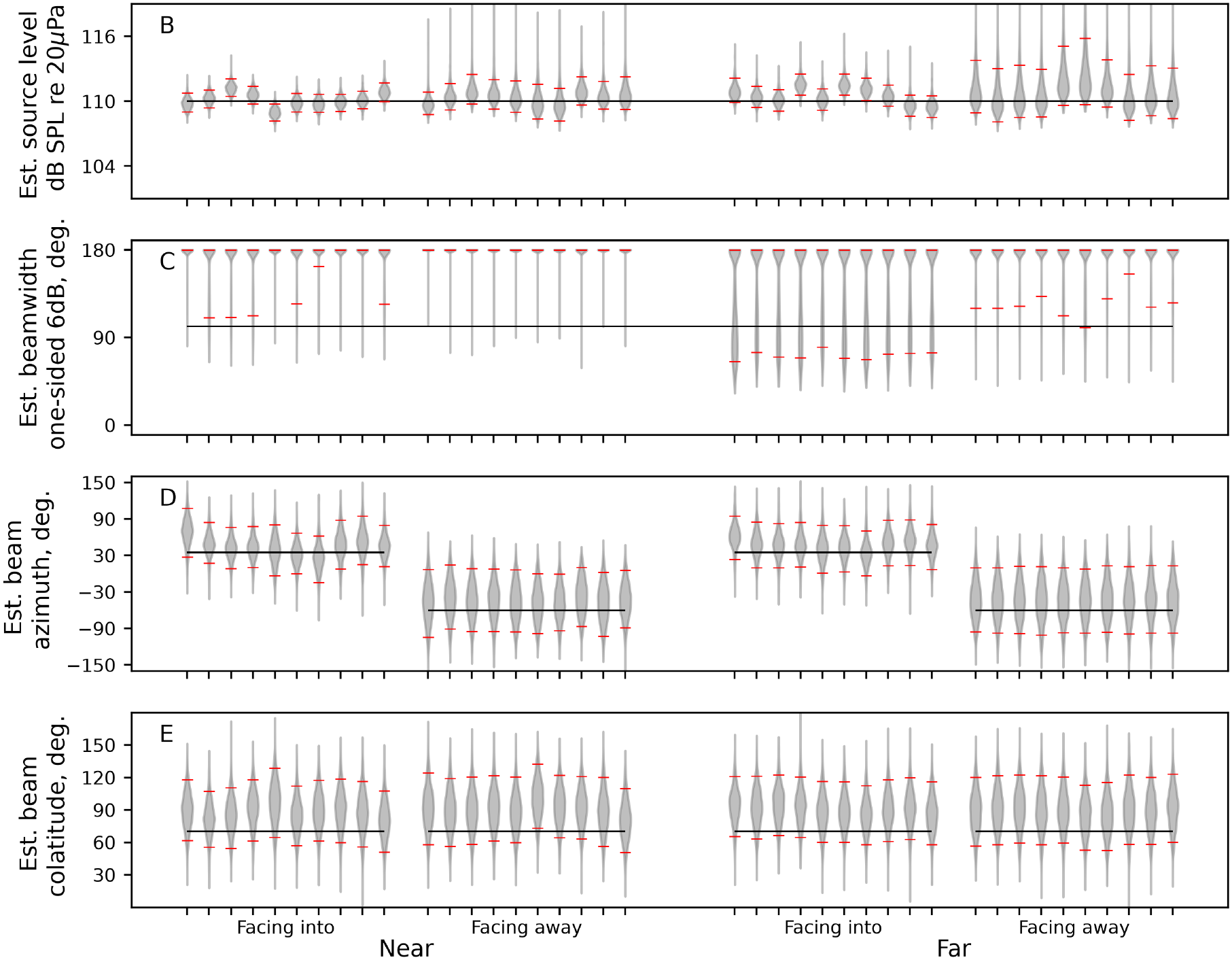
Credible interval coverage of the Giuggioli models (right) for a source at various distances and orientations to the array. Beam aim and source-level can be recovered even when the source is not pointing towards the array, but beam-width cannot be estimated reasonably at all.

**FIG. S4.**
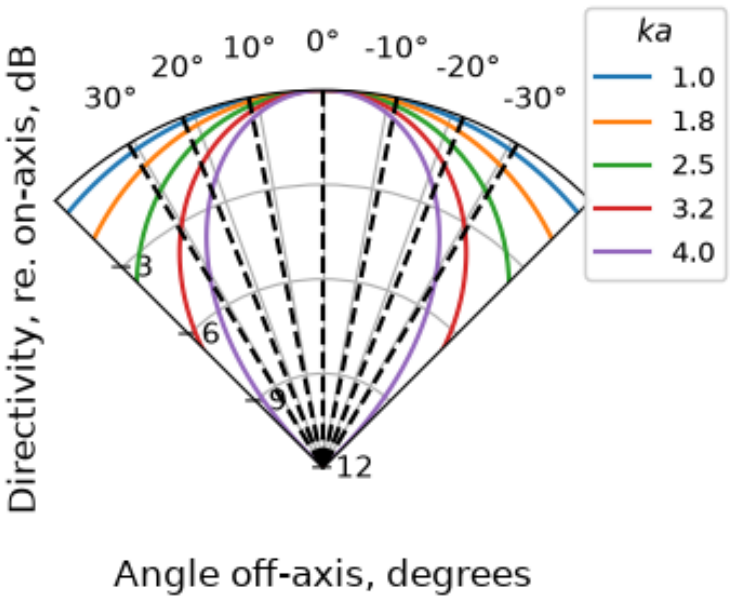
Examining sources of multi-modality in estimating piston in an infinite baffle’s parameters for low ka. The piston model produces very similar predictions for a range of ka values between 1-4 in the first 10 degrees off-axis. When both azimuth and colatitude are considered, 5 of the 12 mics will sample <=10 degrees off-axis - resulting in similar received levels that could correspond to multiple ka values. Black line projections here correspond to the off-axis angle for the simulated source in azimuth. The off-axis colatitude angles will also be the same.

**FIG. S5.**
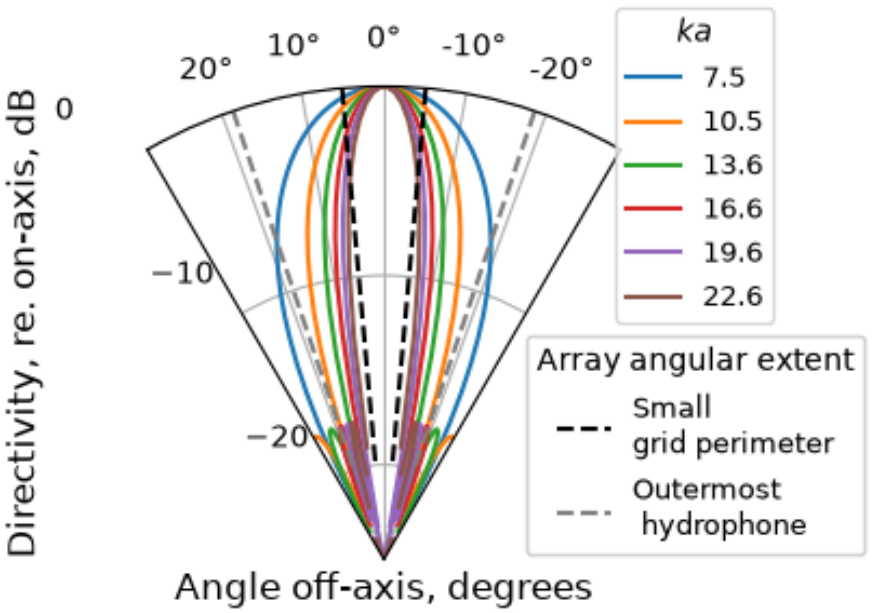
Examining sources of multi-modality in estimating piston in an infinite baffle’s parameters for directional sounds. The piston model is very similar particularly at higher *ka* values, the small grid of 25 hydrophones is also concentrated along the same region that the main lobe of the piston is located (black broken lines indicate the outermost limits). Here 25/48 hydrophones sample a region of the model with very similar values across various ka values.

